# A genome-wide association study highlights a regulatory role for *IFNG-AS1* contributing to cutaneous leishmaniasis in Brazil

**DOI:** 10.1101/2020.01.13.903989

**Authors:** Léa C. Castellucci, Lucas Almeida, Svetlana Cherlin, Michaela Fakiola, Edgar Carvalho, Amanda B. Figueiredo, Clara M. Cavalcanti, Natalia S. Alves, Walderez O. Dutra, Kenneth J. Gollob, Heather J. Cordell, Jenefer M. Blackwell

## Abstract

**Background:** Cutaneous leishmaniasis (CL) caused by *Leishmania braziliensis* remains an important public health problem in Brazil. The goal of this study was to identify genetic risk factors for CL.

**Methods:** Genome-wide analysis was undertaken using DNAs from 956 CL cases and 868 controls (phase 1) and 1110 CL cases and 1178 controls (phase 2) genotyped using Illumina HumanCoreExome BeadChips. Imputation against 1000G data provided 4,498,586 quality-controlled single nucleotide variants (SNVs) common across phase 1 and phase 2 samples. Linear mixed models in FastLMM were used to take account of genetic diversity/ethnicity/admixture. Cellular cytokines were measured using flow cytometry.

**Results:** Combined analysis across cohorts found no associations that achieved genome-wide significance, commonly accepted as *P*<5×10^-8^. Support for variants at wound-healing genes previously studied as candidate genes for CL included *SMAD2* (rs115582038/rs75753347; *P*_imputed_1000G_=1.47×10^-4^). Top novel GWAS hits at P<5×10^-5^ in plausible candidate genes for CL included *SERPINB10* (rs62097497;*P*_imputed_1000G_=2.67×10^-6^), *CRLF3* (rs75270613; *P*_imputed_1000G_=5.12×10^-6^), *STX7* (rs144488134;*P*_imputed_1000G_=6.06×10^-6^), *KRT80* (rs10783496 *P*_imputed_1000G_=6.58×10^-6^), *LAMP3* (rs74285558;*P*_imputed_1000G_=6.54×10^-6^) and *IFNG-AS1* (rs4913269;*P*_imputed_1000G_=1.32×10^-5^). Of these, *LAMP3* (P_adjusted_=9.25×10^-12^; +6-fold), *STX7* (P_adjusted_=7.62×10^-3^; +1.3-fold) and *CRLF3* (P_adjusted_=9.19×10^-9^; +1.97-fold) were all expressed more highly in CL biopsies compared to normal skin, whereas expression of *KRT80* (P_adjusted_=3.07×10^-8^; −3-fold) was lower. Notably, the percent peripheral blood CD3^+^ T cells making interferon-γ in response to *Leishmania* antigen differed significantly by *IFNG-AS1* genotype.

**Conclusions:** In addition to supporting variants in wound-healing genes as genetic risk factors for CL, our GWAS results provide important novel leads to understanding pathogenesis of CL including through the regulation of interferon-γ responses.

## INTRODUCTION

American cutaneous leishmaniasis (ACL) caused by *Leishmania braziliensis* is associated with a variety of clinical presentations including cutaneous leishmaniasis (CL), mucosal leishmaniasis (ML), and disseminated leishmaniasis (DL). ML and DL are generally preceded by CL. CL is the most common form of disease and is associated with localized skin lesions, mainly ulcers, on exposed parts of the body. Whilst normally self-limiting, the degree of pathology and the rate of healing varies, and the lesions leave life-long scars. Not all individuals infected with *Leishmania* parasites go on to develop clinical disease. Subclinical infection is usually associated with a *Leishmania*-specific cellular immune response, measured traditionally as a positive delayed type hypersensitivity or Montenegro skin test responses *in vivo* [1, 2]. Stimulation of peripheral blood lymphocytes with *Leishmania* antigen also results in production of interferon-γ (IFN-γ) and tumour necrosis factor (TNF) in individuals with subclinical infection, but at a lower level than observed with clinical CL [2]. In a longitudinal cohort study, IFN-γ production, mainly by natural killer cells, was associated with protection, but a positive Leishmania-specific skin-test response did not prevent development of disease [3]. Indeed, a positive Leishmania-specific skin-test response has high sensitivity for the diagnosis of CL caused by *L. braziliensis* [1, 2, 4]. Immunologically, the different forms of ACL are generally associated with exaggerated cellular immune responses. In CL, there is a direct correlation between the frequency of CD4^+^ T cells expressing IFNγ and TNF and the size of ulcer [5], while levels of these pro-inflammatory cytokines are higher in ML than CL [6]. The outcome of *L. braziliensis* infection appears to be determined by a fine balance between the pro-inflammatory cytokines IFNγ and TNF and anti-inflammatory interleukin-10 (IL-10) [4, 6, 7].

A major interest has been in determining the extent to which host genetic factors determine these responses. Racial differences, familial clustering and studies in mice all support host genetic control of *Leishmania* infections (reviewed [8–11]). Human family-based genetic epidemiology studies of CL disease caused by the related *L. peruviana* pathogen were consistent with a gene by environment multifactorial model, with a two-locus model of inheritance providing the best fit to the data [12]. This suggested that major genetic risk factors might readily be found for CL. To date, candidate gene studies [13–21] undertaken in *L. braziliensis* endemic regions have demonstrated putative roles for polymorphisms at multiple genes associated with pro- and anti-inflammatory responses (*TNFA*, *SLC11A1*, *CXCR1*, *IL6*, *IL10, CCL2/MCP1*) and with wound healing (*FLI1*, *CTGF*, *TGFBR2*, *SMAD2*, *SMAD3*, *SMAD7*, *COL1A1*) in determining susceptibility to CL or ML disease. This includes susceptibility genes identified from murine studies of leishmaniasis (*SLC11A1*, *FLI1*) [15, 16]. Although frequently underpinned by functional data [17–19] and/or supported by prior immunological studies [22–26], these studies have generally lacked statistical power.

Here we set out to perform a well-powered genome-wide association study for CL caused by *L. braziliensis*. A robust approach incorporating discovery and replication phases was employed using a total sample size across two cohorts that exceeded 2000 CL cases and 2000 controls. No genes/regions achieved genome-wide significance (commonly accepted as P<5×10^-8^ [27–29]) in a combined analysis. Support was found for the prior hypothesis [14, 21] that variants at wound-healing genes are risk factors for CL, including at *SMAD2, SMAD3* and *SMAD 7* studied previously and novel associations at *SMAD1*, *SMAD*4, *SMAD6*, *SMAD9*, *TGFBR3*, *COL11A1* and *COL24A1*. Top novel GWAS hits at P<5×10^-5^ that provide plausible candidate genetic risk factors for CL included variants at *SERPINB10*, *CRLF3*, *STX7*, *LAMP3*, *KRT80* and *IFNG-AS1*. Notable amongst these is the association at *IFNG-AS1* encoding IFNG antisense RNA 1, a role for which is further supported here by functional data showing that the percent of peripheral blood CD3^+^ T cells making IFN-γ or TNF in response to *Leishmania* antigen stimulation differs significantly by *IFNG-AS1* genotype. Given the important role of this long non-coding RNA (IncRNA) in determining expression of the gene *IFNG* encoding IFN-γ [30], our results provide an important novel lead to understanding pathogenesis of CL through the regulation of IFN-γ responses.

## METHODS

### Ethical Considerations, Sampling and Clinical Data Collection

The study was undertaken with approval by the ethical committee of the Hospital Universitário Professor Edgard Santos, Salvador, Brazil (numbers 018/2008 and 22/2012) and the Brazilian National Ethical Committee (CONEP – 305/2007 and CONEP – 1258513.1.000.5537). The enrolment of human subjects complies with the principles laid down in the Helsinki declaration. All participants or their parents/guardians signed written consents to participate in the study and for the storage of de-identified genotype data. Post-quality control (QC) (carried forward below) genotype data will be lodged in the European Genome-phenome Archive upon publication with access controlled through a study-specific Data Access Committee. CL cases and endemic controls were collected from a region of rural rain forest, Corte de Pedra, Bahia, Brazil, where *L. braziliensis* is endemic. Sample collection was based on ascertainment of cases of CL at the Corte de Pedra Public Health Post. Endemic controls were recruited from attendants of cases at the clinic and/or by active follow-up in the field. Collection of cases and controls was undertaken in two phases: phase 1 between March 2008 and April 2010; phase 2 between January 2016 and July 2017. Blood bank controls were volunteer blood bank donors at the HEMOBA Foundation in the city of Salvador collected between the years 2015 and 2017. Basic demographic data (age, sex) were recorded for all participants. Blood (8 ml) was taken by venipuncture and collected into dodecyl citrate acid (DCA)-containing vacutainers (Becton Dickinson). Genomic DNA was prepared using the proteinase K and salting-out method [31]. DNAs were shipped to UK for genotyping at Cambridge Genomic Services, Department of Pathology, University of Cambridge, UK.

### Array Genotyping and Marker QC

DNAs were genotyped on Illumina Infinium® HumanCoreExome Beadchip (Illumina Inc., San Diego, CA, USA), which includes probes for 551,004 single nucleotide variants (SNVs), 282,373 of which are genome-wide tag SNVs that represent core content and are highly informative across ancestries, and 268,631 SNVs that are exome-focused markers. All genotyping data and reference panels were analysed using human genome build 37 (hg19). Individuals were excluded if they had a missing data rate >5%. SNVs were excluded if they had genotype missingness >5%, minor allele frequency (MAF) <0.01, or if they deviated from Hardy-Weinberg equilibrium (HWE; threshold of *P*<1.0×10^-8^). This provided a post-QC dataset of 312,503 genotyped SNVs for the 956 CL cases and 868 controls in the phase 1 sample, and 298,919 genotyped SNVs for the 1110 CL cases and 1178 controls in the phase 2 sample. The phase 1 and phase 2 samples had 52% and 81% power, respectively, and the combined sample 99% power, to detect genome wide significance (*P*<5×10^-8^) for genetic effects with a disease allele frequency of 0.25, effect size (genotype relative risk) of 1.5, and assuming a disease prevalence of 2%.

### SNV Imputation and GWAS

Imputation of missing and non-assayed genetic variants was performed using the 1000 Genomes Project phase 3 reference panel [32], which contains 84.8 million variants for 2504 samples from 26 populations throughout Africa, America, East Asia, Europe and South-East Asia. The 293,563 post-QC array variants genotyped in common across phase 1 and phase 2 samples were phased and imputed using the Michigan Imputation Server v1.0.4 [33]. We excluded imputed SNVs with an information metric <0.8 or genotype probability <0.9, and the remaining variants were converted to genotype calls and filtered for <5% missingness and MAF>0.005. Imputation accuracy was assessed using the r^2^ metric (r^2^>0.5), which represents the squared Pearson correlation between the imputed SNV dosage and the known allele dosage.

Genome-wide association analysis for the CL phenotype was performed using a linear mixed model as implemented in FaST-LMM v2.07, which takes account of both relatedness and population substructure and generally assumes an additive model of inheritance [34]. Population structure and relatedness were controlled using the genetic similarity matrix, computed from 32,696 Phase 1 and 45,569 Phase 2 LD-pruned array variants, and any systematic confounding assessed using quantile-quantile (Q-Q) plots and a test statistic inflation factor (denoted λ) calculated in R version 2.15.0 by dividing the median of the observed chi-squared statistics by the median of the theoretical chi-squared distribution. Genome-wide significance was set at *P*≤5×10^-8^ [27–29]. Manhattan plots were generated using the mhtplot() function of ‘gap’, a genetic analysis package used in R version 3.4.3. Regional plots of association were created using LocusZoom [35] in which -log_10_(P-values) were graphed against their chromosomal location. The 32,696 Phase 1 and 45,569 Phase 2 LD-pruned array variants were matched to data from HapMap populations and PCA plots prepared in R.

### Expression analysis

We examined RNA expression for genes of interest using data from a published microarray study [36] that compared skin biopsies from CL patients from our study area with biopsies of normal skin from non-endemic unexposed donors (accessed from the GEO database: GSE55664). Between group comparisons were made on log transformed data using the GEO2R tool within the GEO database, with P-values adjusted using the Benjamini and Hochberg false discovery rate.

### Antigen-stimulated T cell responses

Peripheral blood mononuclear cells were available for a subset of CL patients used in the present study affording us the opportunity to determine whether antigen stimulated T cell responses differed by *IFNG-AS1* genotype. PBMC from previously untreated cutaneous leishmaniasis patients were obtained from heparinized blood by centrifugation over Ficoll. Cells were stimulated at 1×10^6^ cells/ml with 10 μg/mL soluble Leishmania antigen (SLA) plus 1 μg/mL purified NA/LE anti-human CD28 (clone CD28.2, BD Biosciences, San Jose, CA, USA) at 37°C / 5% CO2 for 15 h and subsequently incubated in the presence of brefeldin A (BD Biosciences) at 37°C with 5% CO2 for up to 4 h. Cells were washed and resuspended in PBS/0.2% BSA for incubation with BUV661 anti-human CD3 monoclonal antibody (UCHT1 clone, BD Biosciences) at 4°C for 30 min in the dark. Cells were fixed, washed, and permeabilized using BD Cytofix/Cytoperm kit before incubation with BV605 anti-human IFN-gamma (B27 clone) and BUV395 anti-human TNF-alfa (MAB11 clone, BD Biosciences) in permeabilization buffer at 4°C for 30 min in the dark. Live and dead cells were distinguished using Fixable Viability Stain 575V (BD Biosciences). The samples were acquired in BD FACSymphony A5 flow cytometer and data analysis was performed using FlowJo 10.6.1 software (BD Biosciences). Differences in responses between genotypes were evaluated in GraphPad Prism using non-parametric Kruskal-Wallis ANOVA with multiple comparisons.

## RESULTS

### Characteristics of the Study Population

Basic demographic information (age, sex) for study participants is shown in Table S1. Cases and controls were well-matched for age and sex overall, the younger age of endemic controls counterbalanced by the older age of blood bank controls. Given ethnic admixture of Brazilian populations it was of interest initially to compare genetic heterogeneity between CL cases and controls, and between the two sources of endemic versus blood bank control samples. PCA plots (Figure 1) show that blood bank controls were generally well matched to endemic controls, with all controls well-matched to cases. There were a small number of outliers in the discovery endemic controls, which also showed greater genetic heterogeneity in the replication sample compared to blood bank controls and cases. Comparison against HapMap populations (Figure 2) showed predominant admixture between Caucasian and African ethnicities in both phase 1 (Figure 2A) and phase 2 (Figure 2B) samples. Linear mixed models were used in association analyses to take account of genetic heterogeneity.

**Figure 1.**
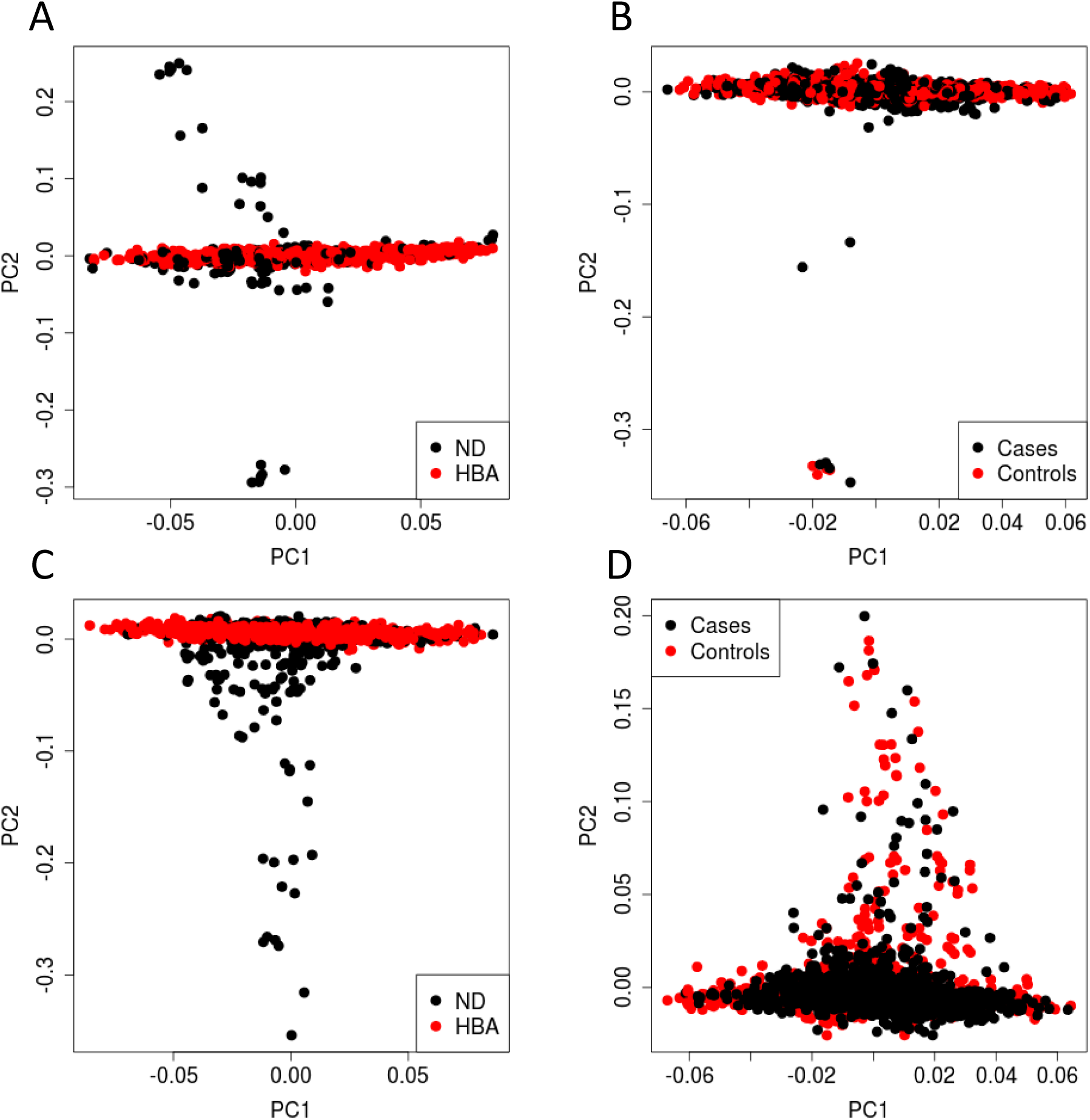
PCA plots comparing genetic heterogeneity across cases and controls used in the study. (A) and (C) compare endemic control (ND) against blood bank controls (HBA) for Phase 1 and Phase 2 samples, respectively. (B) and (D) compare all cases against all controls for Phase 1 and Phase 2 samples, respectively.

**Figure 2.**
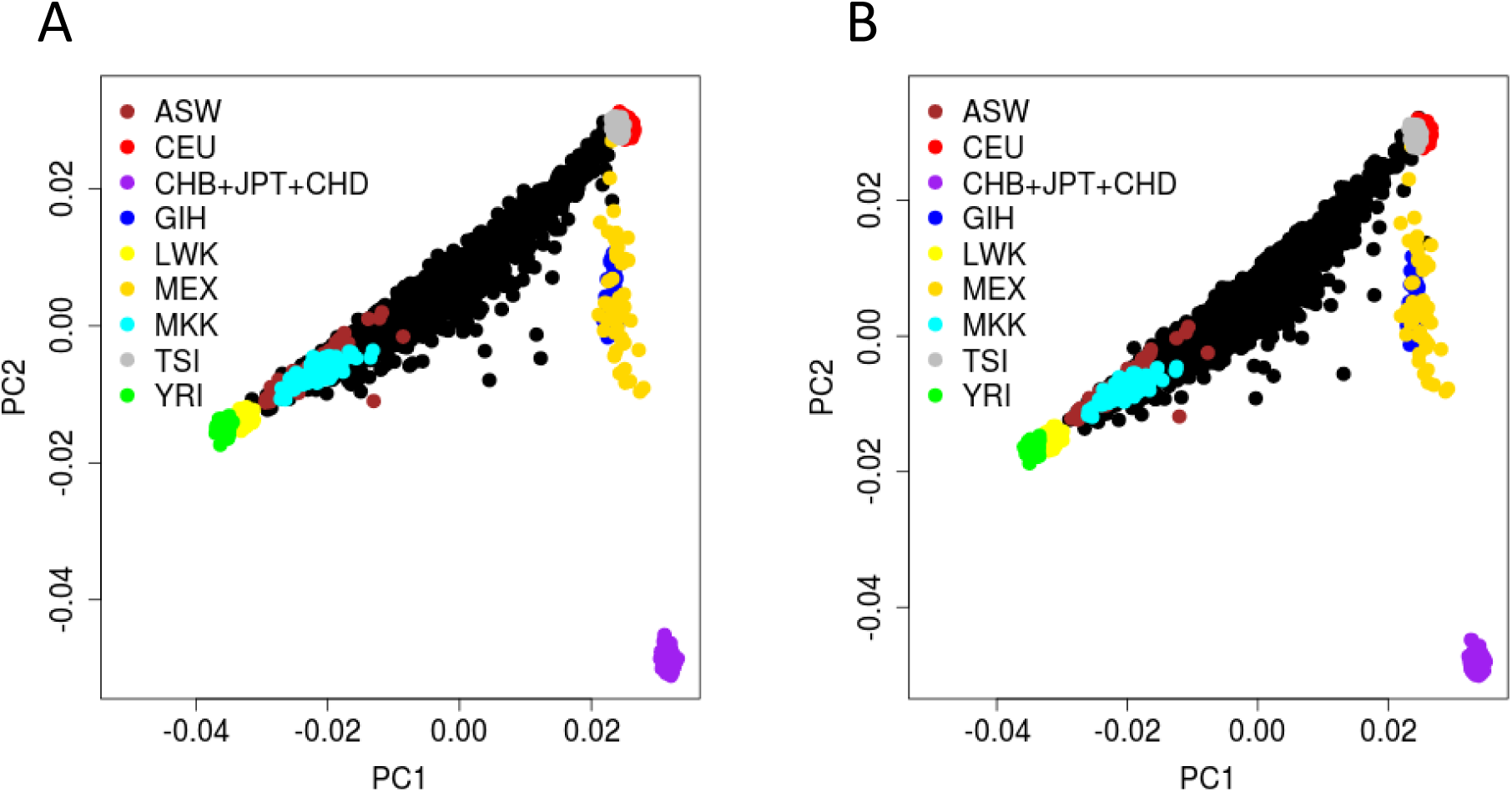
PCA plots comparing genetic heterogeneity in the study sample against HapMap control populations. (A) shows the plot for all Phase 1 samples; (B) shows the plot for all Phase 2 samples. Abbreviations for HapMap populations: ASW - Americans of African Ancestry in SW US; CEU - Utah Residents (CEPH) with Northern and Western European Ancestry; CHB - Han Chinese in Beijing, China; JPT - Japanese in Tokyo, Japan; CHD - Chinese in Metropolitan Denver, Colorado; GIH - Gujarati Indian from Houston, Texas; LWK - Luhya in Webuye, Kenya; MEX - Mexican ancestry in Los Angeles, California; MKK - Maasai in Kinyawa, Kenya; TSI - Toscani in Italia; YRI - Yoruba in Ibadan, Nigeria.

**Table 1.** Top GWAS Hits in Genes of Plausible Functional Interest as Genetic Risk Factors for CL Caused by *L. braziliensis*.

| Chr | Position (bp) | rsID | P-value | Odds Ratio (95% CI) | Beta (SE) | Allele<sup>1</sup> | Variant Origin | Location | Gene | Function |
| --- | --- | --- | --- | --- | --- | --- | --- | --- | --- | --- |
| 3 | 182857261 | rs74285558 | 6.54E-06 | 0.87 (0.82-0.92) | -0.034 (0.008) | A (C/A) | Global | intron | LAMP3 | Lysosomal associated membrane protein 3 |
| 6 | 132815562 | rs144488134 | 6.10E-06 | 0.82 (0.75-0.89) | -0.034 (0.007) | A (C/A) | African | intron | STX7 | Syntaxin 7 |
| 12 | 52590004 | rs10783496 | 6.58E-06 | 1.06 (1.03-1.09) | 0.035 (0.008) | A (G/A) | Global | intron | KRT80 | Keratin 80 |
| 12 | 68407845 | rs4913269 | 1.32E-05 | 1.06 (1.03-1.08) | 0.033 (0.008) | G (C/G) | Global | intron | IFNG-AS1 | IFNG antisense RNA 1 |
| 17 | 29136126 | rs75270613 | 5.12E-06 | 0.83 (0.77-0.90) | -0.034 (0.008) | A (C/A) | African | intron | CRLF3 | Cytokine receptor like factor 3 |
| 18 | 61598763 | rs8084306 | 1.56E-06 | 1.07 (1.04-1.10) | 0.038 (0.008) | C (T/C) | Global | intron | SERPINB10 | Serpin family B member 10 |

**NOTE** Details of all top hits at $P < 5 \times 10^{-5}$ are provided in Table S1. <sup>1</sup>Associated allele (ancestral/minor) for risk or protection as indicated by the odds ratio.

### Genome-wide Association Study

The GWAS was performed in two phases. Manhattan plots for genotyped data for phase 1 and phase 2 are shown in Figure S1. Quantile-quantile plots (Figure S2) of the observed versus expected -log_10_ P-values did not show any evidence for systematic bias in either phase 1 or phase 2, with inflation factors λ of 0.998 and 1, respectively. A Manhattan plot for the combined analysis of imputed genotype data is shown in Figure 3. The top hits did not achieve genome-wide significance, commonly accepted as P<5×10^-8^ for GWAS using variants with minor allele frequencies >1% [27–29].

**Figure 3.**
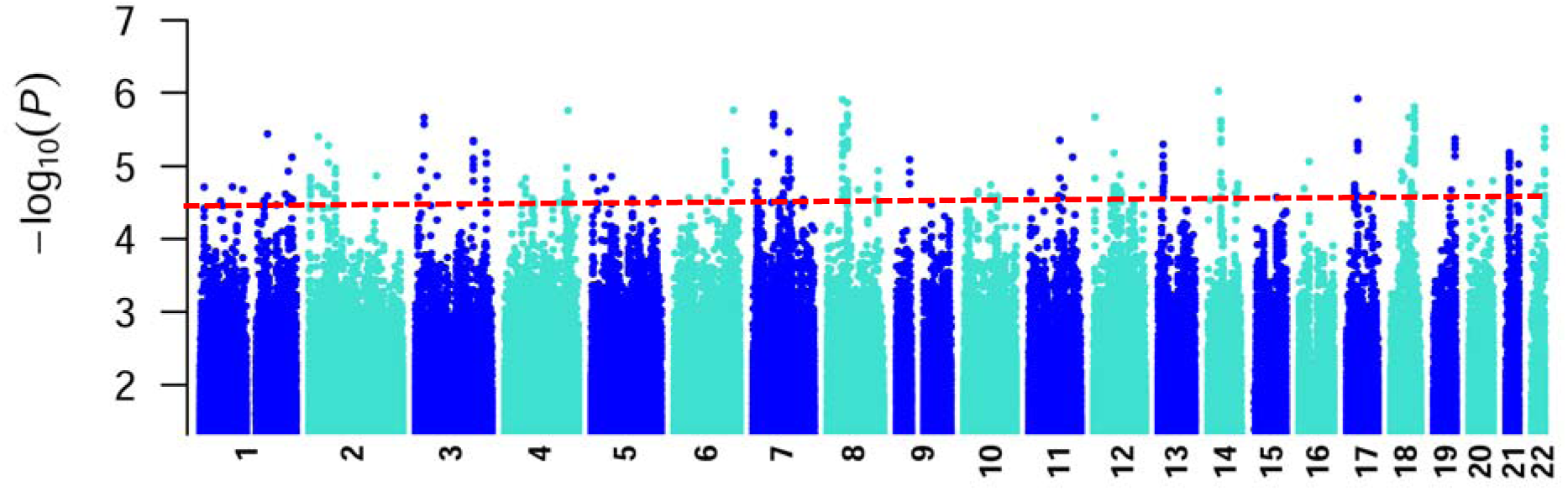
Manhattan plot of results from the combined analysis for the 4.46M high-quality 1000G imputed SNV variants common to Phase 1 and Phase 2 samples. Data are for analysis in FastLMM looking for association between SNVs and CL. The Y-axis indicates –log10 *P* values for association, the X axis indicates the positions across each chromosome.

### Interrogating the GWAS Data in Relation to Previous Candidate Gene Studies

As noted above, previous candidate gene studies attempting to identify genetic risk factors for CL caused by *L. braziliensis* have typically been small, underpowered family-based or case-control studies [13–21]. We therefore interrogated the GWAS data to determine whether evidence could be found to support the candidacy of these genes. Table S2 presents the top hits for previously studied candidate, or functionally related, genes. Given *a priori* evidence to look at these genes, we used P<0.01 as a cut-off to identify possible associations. No variants were associated at P<0.01 for *TNFA*, *SLC11A1*, *CXCR1*, *IL6*, *IL10*, *CCL2/MCP1*, *FLI1*, *CTGF*, *COL1A1*, or *TGFBR2*. Associations were observed for variants at *SMAD2*, *SMAD3* and *SMAD7* (Table S2), wound healing genes that had previously been shown to be associated with CL [14]. The strongest signal was at *SMAD2*, as indicated by the peak of association at multiple variants observed on the LocusZoom plot across the region (Figure 4A). Associations at *SMAD3* and *SMAD7* (Table S2; see also Figure 4A LocusZoom plot that includes *SMAD7*) were less well-supported. Some evidence for associations at related SMADs was also observed (Table S2), specifically at *SMAD1*, *SMAD4* (see also Figure 4A), *SMAD6* and *SMAD9*. The major role for SMAD proteins is to transduce signals from receptors of the transforming growth factor beta (TGFβ) superfamily. Although no association at P<0.01 was observed at *TGFBR2*, association (Table S2) at the functionally related gene *TGFBR3* (Figure 4B) was supported at multiple variants across the gene. Similarly, associations (Table S2) observed at collagen genes, *COL24A1* (Figure 4C) and *COL11A1* (Figure 4D), functionally related to wound healing gene *COL1A1* were supported at multiple variants across each gene. Associations were also observed (Table S2) for variants at genes (*IL6R* and *IL10R*) encoding receptors for cytokines IL-6 and IL-10 that have previously been shown to be genetically and/or functionally associated with cutaneous forms of disease caused by *L. braziliensis* [17, 18].

**Figure 4.**
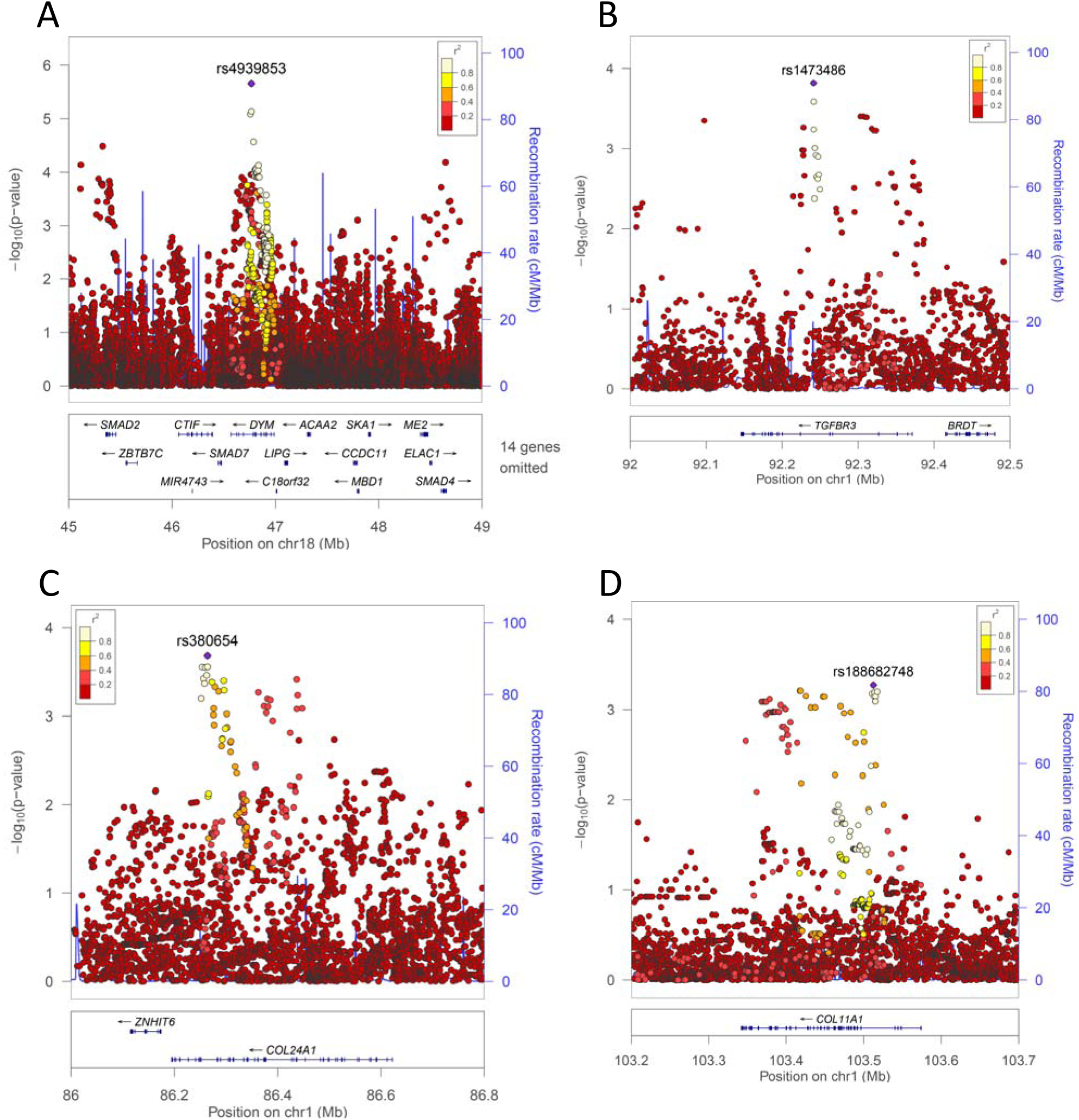
LocusZoom plot of single-nucleotide variant (SNV) associations with CL across regions of chromosomes containing genes previously associated with, or functionally related to, CL in candidate genes studies. The –log10 *P* values (left y-axis) are shown in the top section of the plot. Dots representing individual SNVs are color coded (see key) based on their population-specific linkage disequilibrium r2 with the top SNV (annotated by rs ID) in the region. The right Y-axis is for recombination rate (blue line), based on HapMap data. The bottom section of each plot shows the positions of genes across the region. The genes of interest include (A) *SMAD2*, *SMAD7* and *SMAD4* (for clarity, 14 genes are omitted from the bottom section of this plot; association at *DYM* is listed in Table S1); (B) *TGFBR*3; (C) *COL24A1*; and (D) *COL11A1*.

### Candidate Top Hits from the GWAS Data

Table S3 provides a full list of hits that were suggestive of association, using a cut-off P<5×10^-5^. Table 1, Figure 5 and Figure S3 highlight those hits that are of potential functional relevance to CL. Of specific interest are genes, *LAMP3* (Figure 5A) and *STX7* (Figure 5B), that play a role in lysosome function. *LAMP3* encodes lysosomal associated membrane protein 3, also known as dendritic cell LAMP or DCLAMP. *STX7* encodes Syntaxin 7, which plays a role in the ordered fusion of endosomes and lysosomes with phagosomes. Other genes, *KRT80* (Figure 5C) and *CRLF3* (Figure S1A), may relate to perturbations in skin at the site of infection, while *SERPINB10* (Figure S1B) and *IFNG-AS1* (Figure 5D) are highlighted in relation to more central roles in determining immune responses. Other hits at P<5×10^-5^ are shown in Table S3. Whilst variants in these regions may be true CL susceptibility loci, there is currently no obvious support for their candidacy related to their functions.

**Figure 5.**
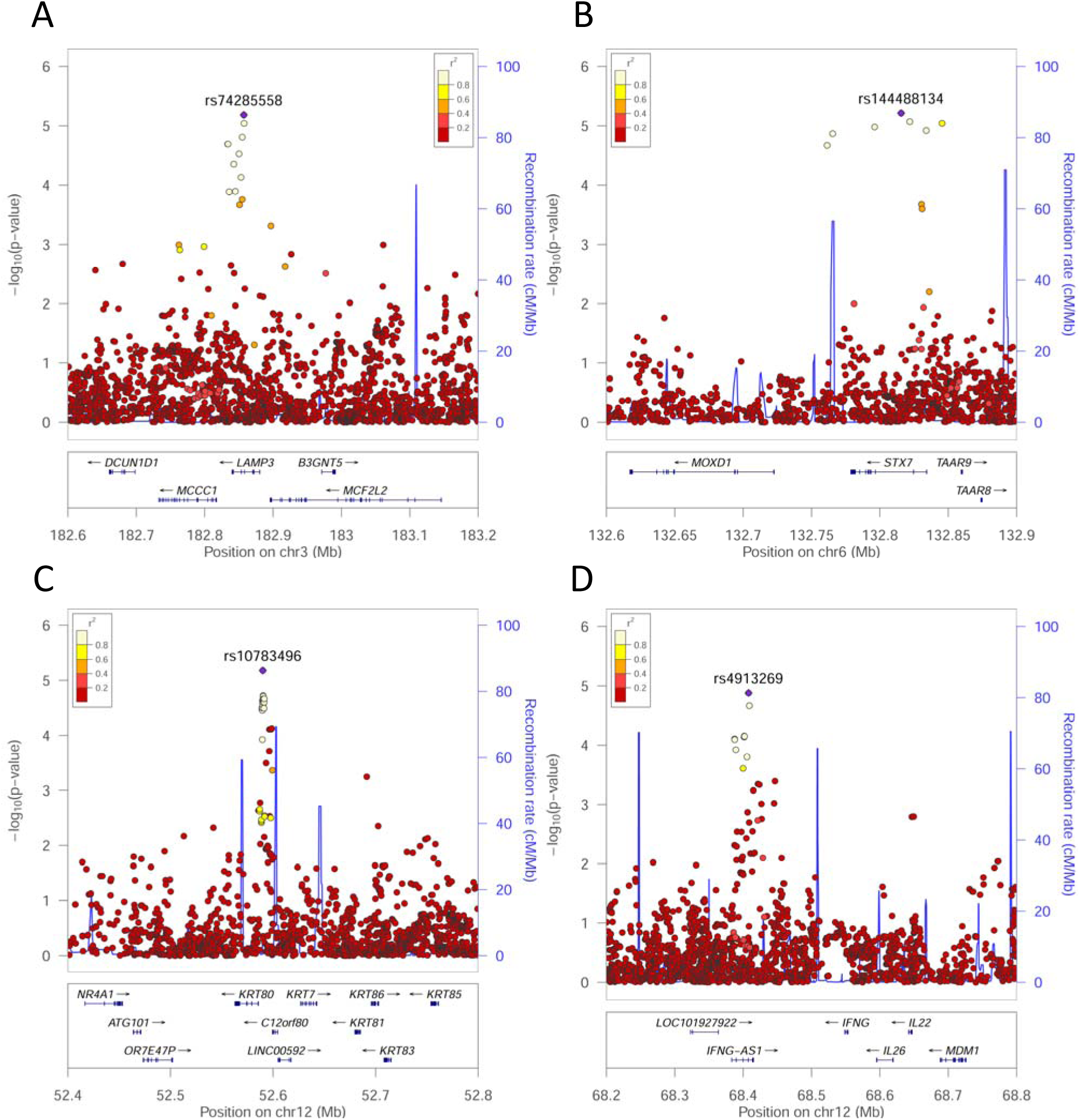
LocusZoom plot of single-nucleotide variant (SNV) associations across regions of chromosomes containing genes associated with CL at P<5×10^-5^ in this study. The –log10 *P* values (left y-axis) are shown in the top section of each plot. Dots representing individual SNVs are color coded (see key) based on their population-specific linkage disequilibrium r2 with the top SNV (annotated by rs ID) in the region. The right Y-axis is for recombination rate (blue line), based on HapMap data. The bottom section of each plot shows the positions of genes across the region. The genes of interest include (A) *LAMP3*; (B) *STX7*; (C) *KRT80*; and (D) *IFNG-AS1*. Plots for two other regions containing genes of specific interest, *CRLF3* and *SERPINB10*, are provided in Figure S3.

### Expression Analysis of Novel Candidate Genes Arising from the GWAS

Disease associated SNVs identified through GWAS mostly localize to non-coding sequence [37] and are enriched for expression quantitative trait loci [38, 39]. An initial piece of evidence to support candidacy of variants identified in our GWAS was to determine (a) if they were expressed in skin and/or lesions, and (b) whether their expression in CL lesions was up- or down-regulated relative to expression in normal skin. We therefore interrogated previously published [36] microarray data that had compared RNA from lesions biopsies with RNA from normal skin biopsies (Figure S4). This showed that *LAMP3* (P_adjusted_ = 9.25×10^-12^; +6-fold increase in lesions compared to normal skin), *STX7* (P_adjusted_ = 7.62×10^-3^; +1.3-fold) and *CRLF3* (P_adjusted_ = 9.19×10^-9^; +1.97-fold) were all up-regulated in CL biopsies, whereas expression of *KRT80* (P_adjusted_ = 3.07×10^-8^; −3-fold) was down-regulated. Individual variation in expression levels of these genes in lesions from different donors suggests that some additional level of regulated gene expression may exist. *SERPINB10* was expressed at very low levels in both normal and lesion biopsy samples. Probes for *IFNG-AS1* were not present on the chips used in the published microarray data [36]. Since IFNG-AS1 expression is known to influence IFN-γ production [30], we therefore sought functional support for this gene by examining the percent of antigen-stimulated T cells producing IFN-γ by *IFNG-AS1* genotypes (Figure 6). A significant difference across genotypes was observed at rs4913269 (ANOVA P_adjusted_=0.044), with individuals homozygous for the disease-associated G allele showing a significantly lower percentage of IFN-γ producing T cells compared to heterozygous individuals (Figure 6A). A similar observation was made for the percentage of T cells producing TNF (Figure 6B; ANOVA P_adjusted_=0.021), which was strongly correlated with the percentage of IFN-γ producing T cells (Figure 6C; r^2^=0.31. P=0.0003). Parallel observations were made for the second top SNP rs11177004 (Figure 6D-F), and for 5 other SNPs in strong linkage disequilibrium (Figure 5D) with these two top hits (data not shown).

**Figure 6.**
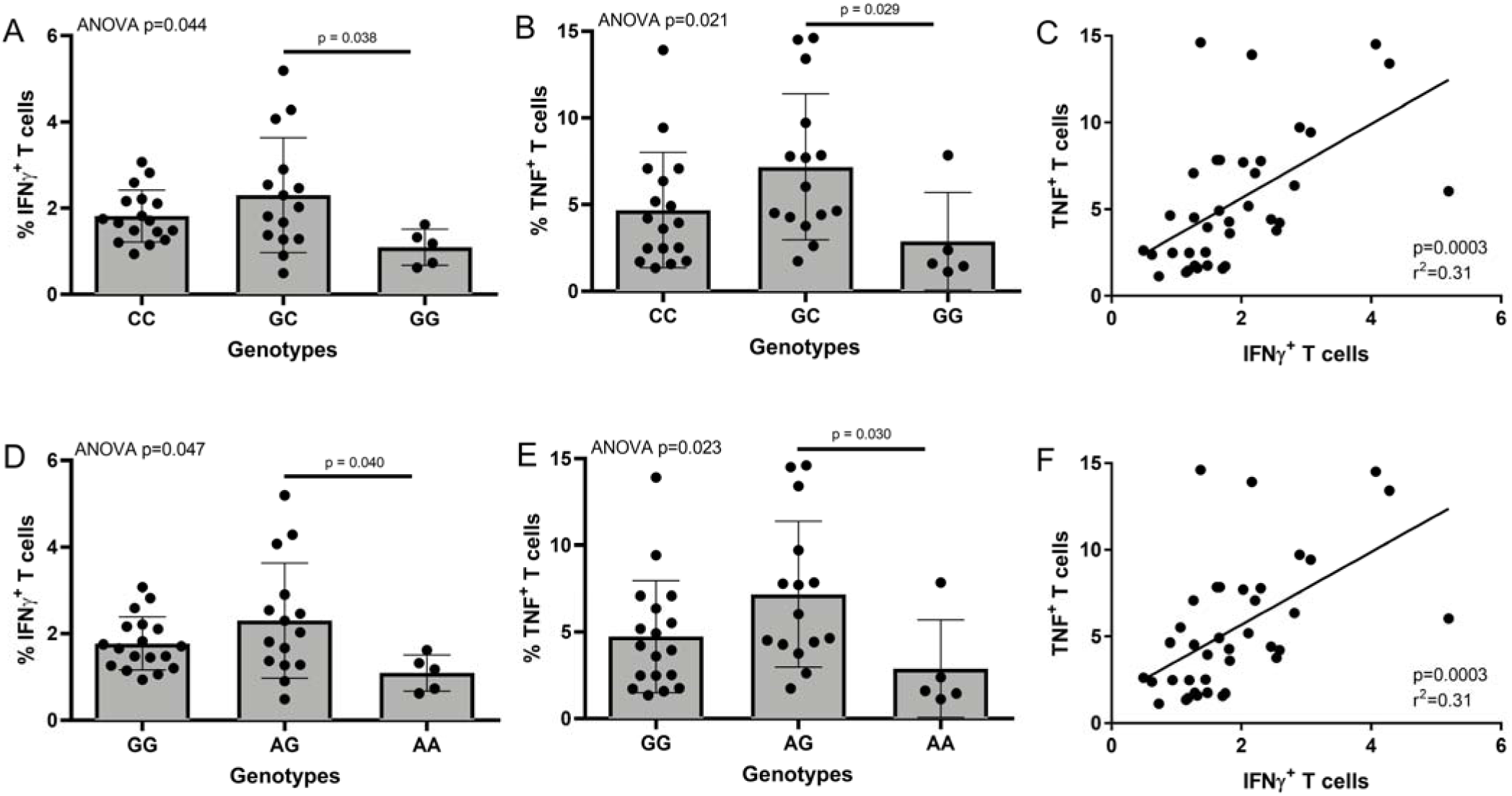
Plots showing differences in percentages of antigen-stimulated IFN-γ and TNF producing CD3+ T cells by *IFNG-AS1* genotype. Plots (A), (B) and (C) show results by genotype for the top SNV rs4913269 at Chromosome 12 bp position 68407845, (D), (E) and (F) for the second top SNV rs11177004 at Chromosome 12 bp position 68409009 (Build 37). (A) and (D) show the percent IFN-γ^+^ CD3^+^ T cells by genotype; (B) and (E) the percent TNF^+^ CD3^+^ T cells by genotype, and (C) and (F) show the correlation between percent IFN-γ^+^ and percent TNF^+^ CD3^+^ T cells for individuals genotyped.

## DISCUSSION

Results of the GWAS presented here provide the first hypothesis-free insights into genetic risk factors for CL caused by *L. braziliensis*. Despite prior evidence for genetic regulation of *Leishmania* infections [8–12], and the well-powered phase 1 and phase 2 study undertaken here, no signals of association achieved genome-wide significance generally accepted as 5×10^-8^ for this type of GWAS [27–29]. Nevertheless, interesting novel genetic risk factors were supported by interrogating suggestive associations that achieved P<5×10^-5^, and modest support (P<0.01) was obtained for some previously studied candidate genes for CL.

In relation to previous candidate gene studies, the concept of a role for wound healing genes in contributing to CL [14, 20] was supported by associations at SMAD genes, with the strongest support observed for variants at *SMAD2*. SMAD proteins transduce signals from receptors of the TGFβ superfamily. Although we did not replicate our earlier study [14] showing a genetic association with the type II receptor for TGFβ encoded by *TGFBR2*, we did see a positive association at *TGFBR3* encoding the type III receptor. TGFBR3 is a membrane proteoglycan [40] that often functions as a co-receptor with other TGFβ receptor superfamily members [41]. Shedding of the ectodomain also produces soluble TGFBR3 which may inhibit TGFβ signalling [40]. TGFβ has long been known to play an important role in tissue fibrosis [42], with abnormal TGFβ regulation and function implicated in a large number (reviewed [43]) of fibrotic and inflammatory pathologies. CL lesions are characterised by a combination of inflammatory immune processes and fibrosis [44]. The fibrotic reaction leads to increased production of extracellular matrix proteins including collagens, as well as the activation of local fibroblasts to differentiate into myofibroblasts [45]. TGFβ stimulates COL11A1 expression in dermal fibroblasts [46], while novel myofibroblast-specific expression of COL24A1 is a signature of skin wounds [47]. Together with associations at *IL6R* and *IL10R*, our GWAS results support the complex interplay between fibrotic and inflammatory processes in determining pathologies associated with CL caused by *L. braziliensis*.

The benefit of using an unbiased genome-wide approach to identification of genetic risk factors for disease is its ability to highlight novel genes/pathways involved in disease pathogenesis. Two genetic associations were identified which related to intracellular localization of *Leishmania* parasite in phagolysosomes [48, 49], namely the associations at *LAMP3* and *STX7*. *LAMP3* encodes lysosomal associated membrane protein 3, also known as dendritic cell LAMP or DCLAMP [50]. Expression of DCLAMP increases markedly upon activation of dendritic cells where it localizes to the MHC Class II compartment immediately before the translocation of MHC Class II molecules to the cell surface [50]. LAMP3 was expressed at much higher levels in RNA from lesion biopsies compared to normal skin, supporting a role for this gene in contributing to disease susceptibility. Since dendritic cells are the most potent antigen-presenting cells that induce primary T-cell responses, it is likely that the influence of variants at *LAMP3* will be related to the processing and presentation of *Leishmania* antigens to T cells. *STX7*, on the other hand, encodes syntaxin 7 which influences vesicle trafficking to lysosomes, including phagosome-lysosome fusion [51]. Variants at this gene are therefore likely to be influencing the delivery of *Leishmania* phagosomes to the lysosomal compartment of host macrophages. STX7 was expressed in normal skin with a modest increase in expression in lesion biopsies, again supporting a possible role for this gene in contributing to disease pathology.

Other genes identified included *KRT80* and *CRLF3* that may relate to perturbations in the skin at the site of infection and/or in the wound healing response. Keratin 80 encoded by *KRT80* is a type II epithelial keratin that is an intermediate filament protein responsible for structural integrity of epithelial cells and has biased expression in skin keratinocytes [52]. At the time of infection pathogens invading the skin can cause keratinocytes to produce chemokines which attract monocytes, natural killer cells, T cells, and dendritic cells [53]. Interestingly, *KRT80* and multiple other keratins (*KRT77*, *KRT81*, *KRT4*, *KRT39*, *KRT32*, *KRT33B*; data not shown) were expressed at much lower levels in lesion biopsies compared to normal skin. To what extent this represents a paucity of keratinocytes in lesions, as opposed to specific down regulation of gene expression in keratinocytes within lesions, will require further investigation. Keratinocytes are known to play an important role in wound healing [45], are potent producers of IL10 and TGFβ [54], and have been shown to change to a sclerotic phenotype by gene silencing of Fli1 [55]. Although association of human CL and *FLI1* [14, 15] was not replicated here, *Fli1* is a confirmed CL susceptibility gene in mice [56]. Hence, our novel associations also continue to focus on molecules and cells involved in wound healing. *CRLF3*, on the other hand, encodes a member of the cytokine receptor-like factor 3 family which is expressed in normal skin, but shows pathologically enhanced expression in premalignant actinic keratosis and malignant squamous cell carcinoma [57]. Further work will be required to determine if CRLF3 is similarly dysregulated in CL lesions.

Two further genes, *SERPINB10* and *IFNG-AS1*, are highlighted in relation to more central roles in determining immune responses. Serpin family B member 10 encoded by *SERPINB10* is a peptidase inhibitor expressed in bone marrow [58] in haematopoietic cells of the monocytic lineage [59] and is known to inhibit TNF-induced apoptosis [60]. Epithelial SERPINB10 contributes to allergic eosinophilic inflammation [61], suggesting additional roles for this molecule in immunopathology. However, we found no evidence for expression of SERPINB10 in normal skin or in lesion biopsies, suggesting that a role in determining susceptibility to CL must be at the level of central regulation of haematopoiesis and/or immune responses. IFNG antisense RNA 1 encoded by *IFNG-AS1* has been shown to fine-tune the magnitude of IFN-γ responses [30]. It is expressed in both mouse and human T helper1 cells and positively regulates *Ifng* expression [62, 63]. Transient over-expression of *Ifng-as1* in mice is associated with increased IFN-γ and reduced susceptibility to *Salmonella enterica* [62]. Conversely, deletion of *Ifng-as1* in mice compromises host defence against *Toxoplasma gondii* infection by reducing *Ifng* expression. Discordant expression of *IFNG* and *IFNG-AS1* is seen in long-lasting memory T cells, where high expression of *IFNG-AS1* was associated with low *IFNG* suggesting that there may be feedback inhibition [30]. Here we observed that *IFNG-AS1* genotype was associated with downstream effects on both the percentage of IFN-γ-producing CD3^+^ T cells and the highly correlated percentage of TNF-producing CD3^+^ T cells following stimulation of peripheral blood mononuclear cells with *Leishmania*-derived antigens. In particular, individuals who were homozygous for the disease-associated allele at each of the top 7 *IFNG-AS1* associated SNVs had significantly lower percentages of IFN-γ and TNF producing T cells, suggesting that lower IFN-γ and TNF production regulated by *IFNG-AS1* is associated with increased disease risk. Our results indicate that regulatory variants at *IFNG-AS1* may act as risk factors for CL by modulating production of these two pro-inflammatory cytokines known to be important in activating macrophages for anti-leishmanial activity.

## CONCLUSIONS

In conclusion, while not achieving genome-wide significance at any loci, our GWAS has provided further support for variants in wound-healing genes as genetic risk factors for CL as well as providing important novel leads to understanding pathogenesis of CL including through the regulation of IFN-γ responses.

## *Acknowledgements*.

We would like to thank the staff of the Health Post at Corte de Pedra for their assistance in the collection of samples and clinical and field data, as well as staff at the HEMOBA Foundation Blood Bank in Salvador.

## Notes

### Disclaimer

The study sponsor had no role in study design, data collection, data analysis, data interpretation, or writing of the report. The corresponding author had full access to all study data and had final responsibility for the decision to submit for publication.

---

**Figure S1.**
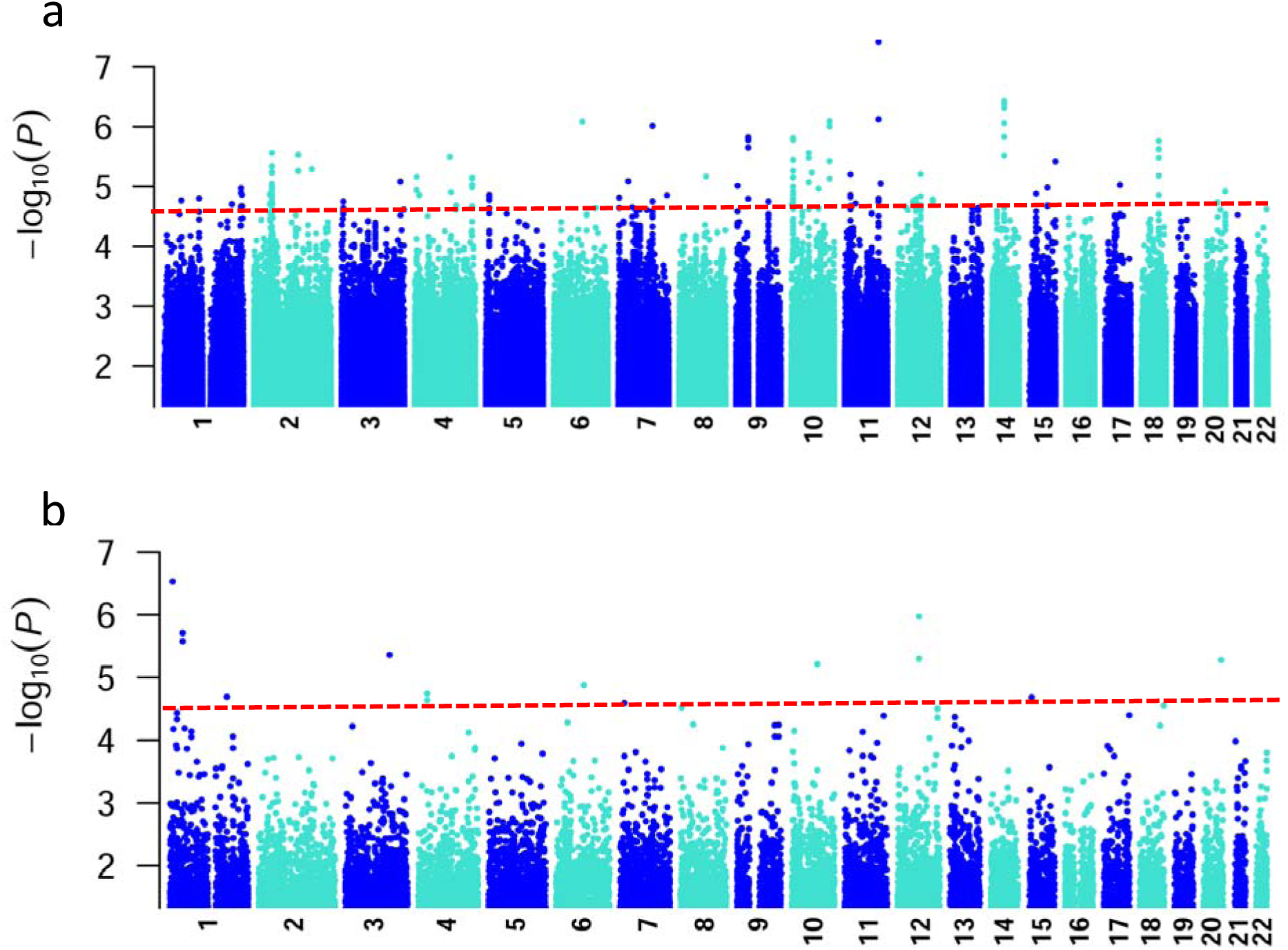
Manhattan plots of results of association analyses for genotyped SNVs. (A) for Phase 1. (B) for Phase 2. Data are for analysis in FastLMM looking for association between SNVs and CL. The Y-axis indicates – log10 *P* values for association, the X axis indicates the positions across each chromosome. The red dotted lines indicate the *P*=5×10^-5^ cut-off used to look for suggestive associations in the combined analysis (main Fig. 3).

**Figure S2.**
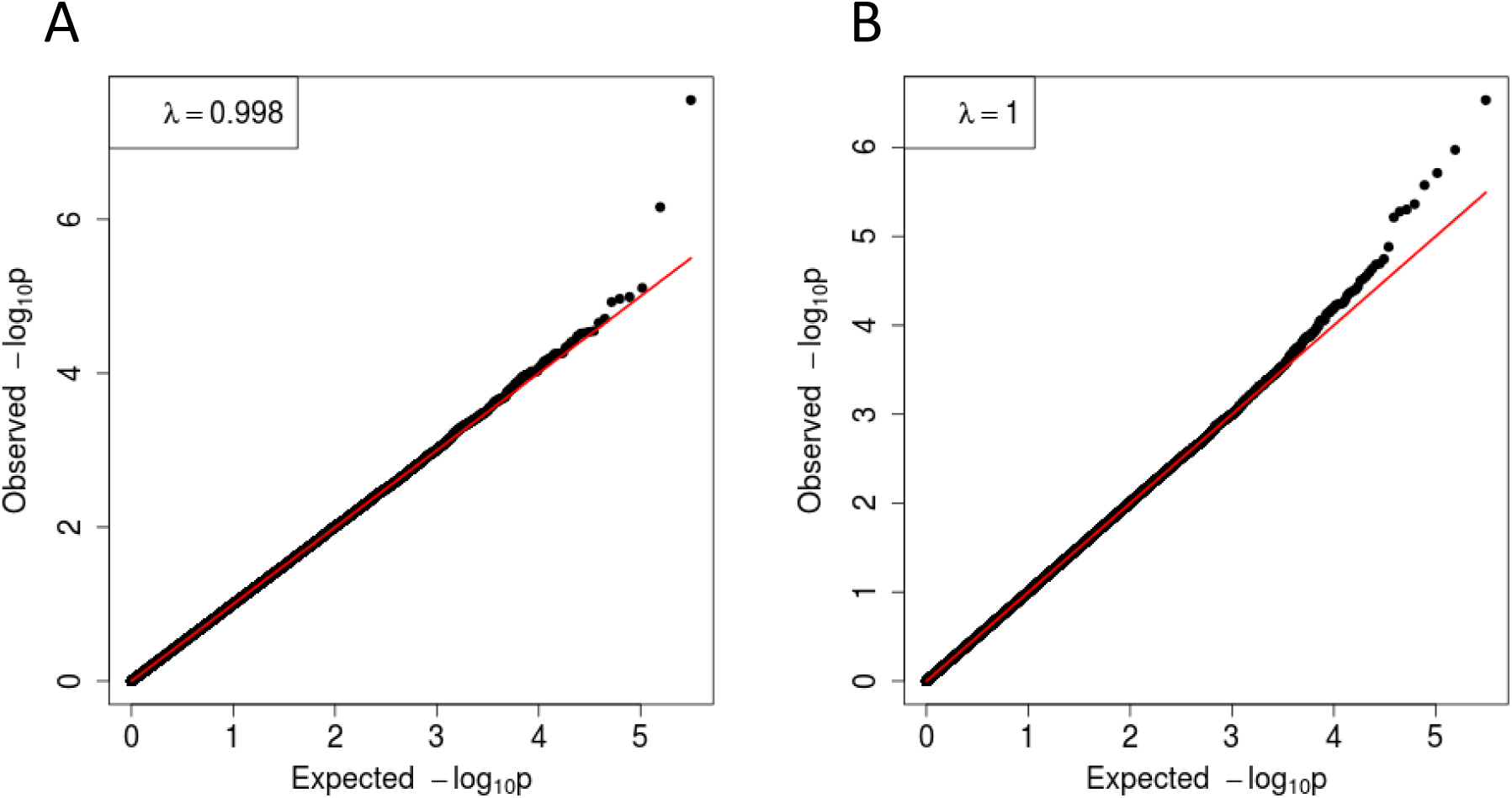
Quantile-quantile plot of GWAS p-values for genotyped data for (A) Phase 1; and (B) Phase 2 of the study.

**Figure S3.**
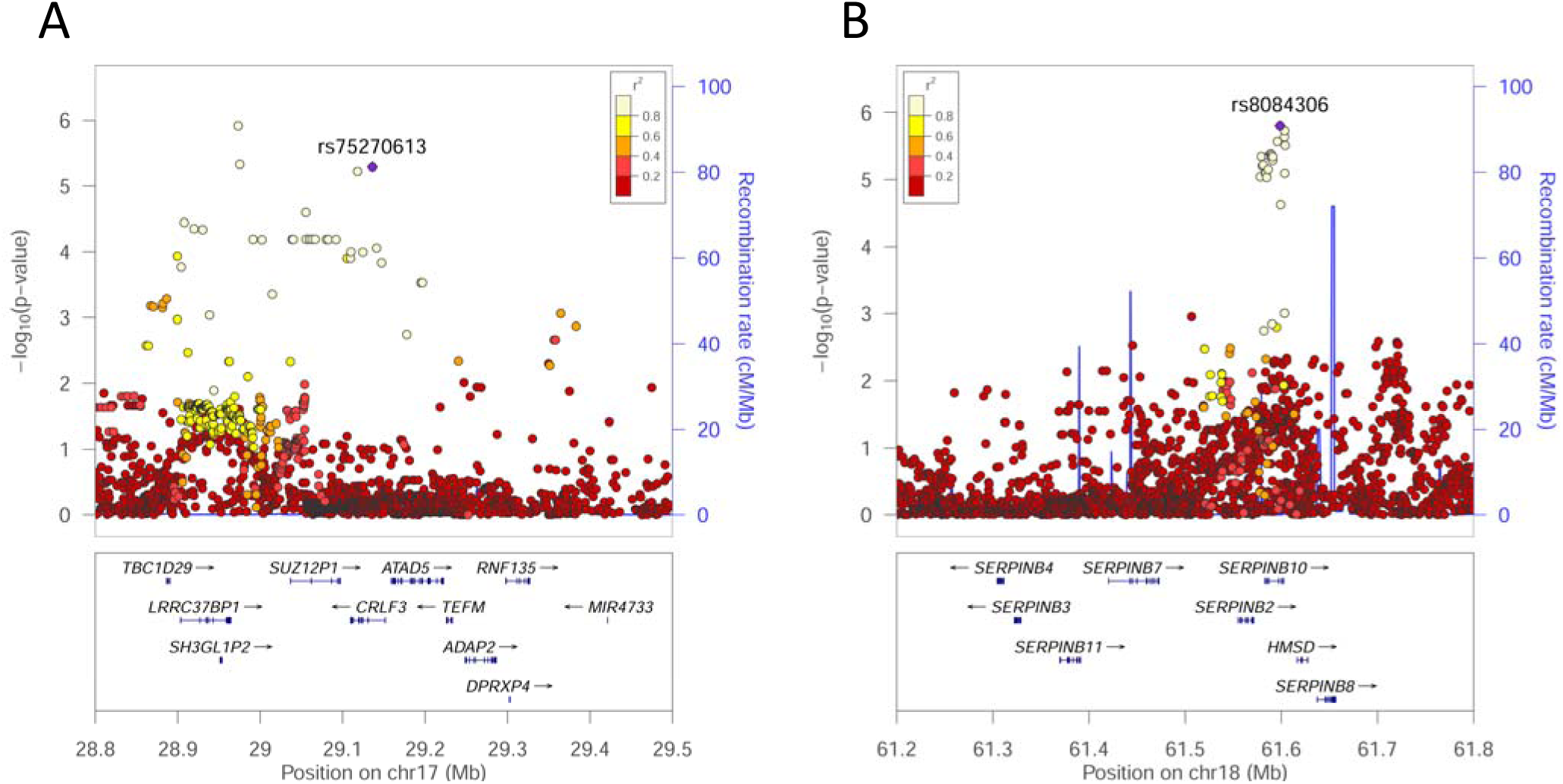
Regional association plots (LocusZoom^1^ of the signal for CL association in the regions of (A) *CRLF3* on Chromosome 17; and (B) *SERPINB10* on Chromosome 18. The −log_10_ P-values are shown on the upper part of each plot. Dots representing individual SNVs are color coded (see key) based on their population-specific linkage disequilibrium r2 with the top SNV (annotated by rs ID) in the region. The right Y-axis is for recombination rate (blue line), based on HapMap data. The bottom section of each plot shows the positions of genes across the region.

**Figure S4.**
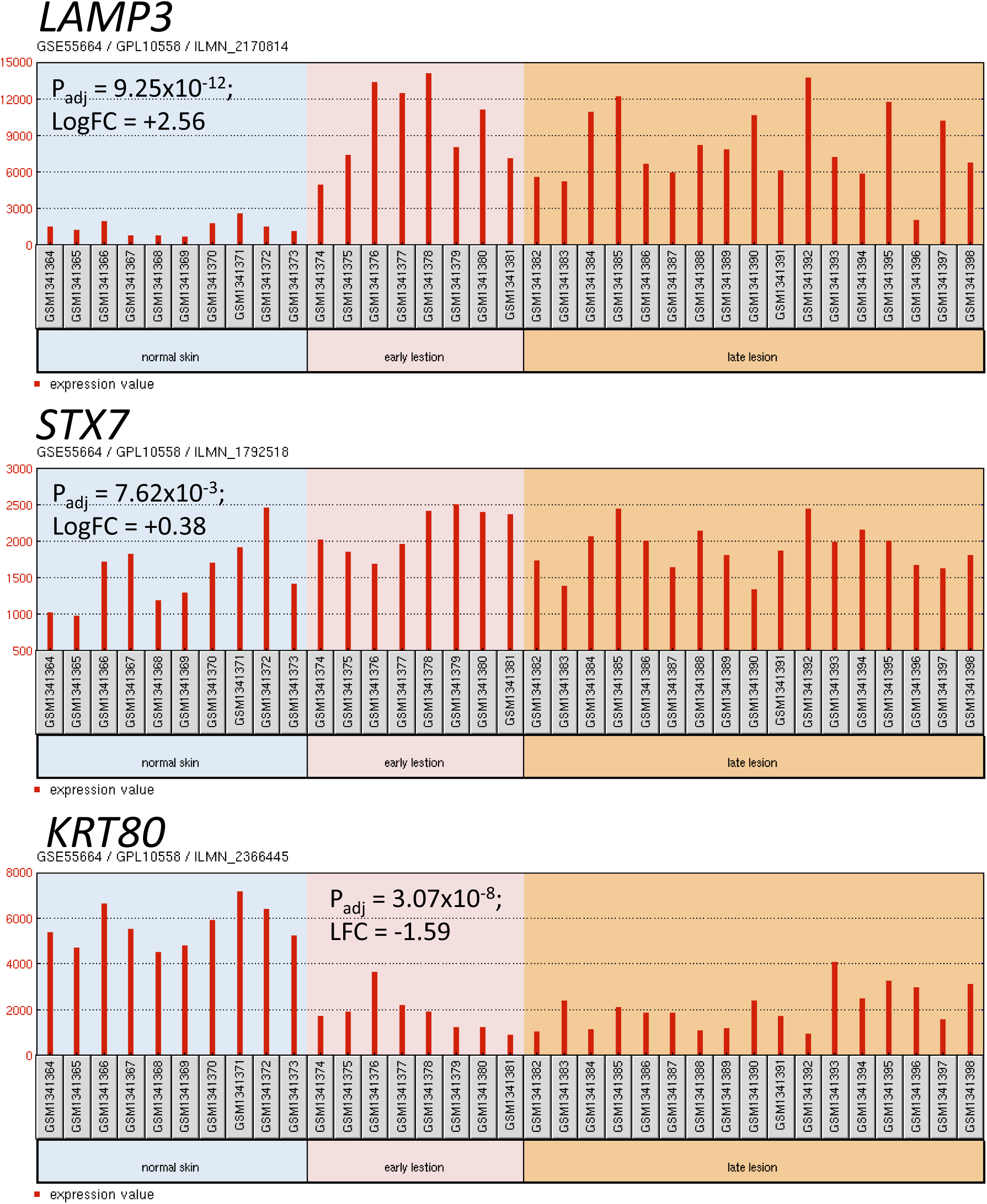
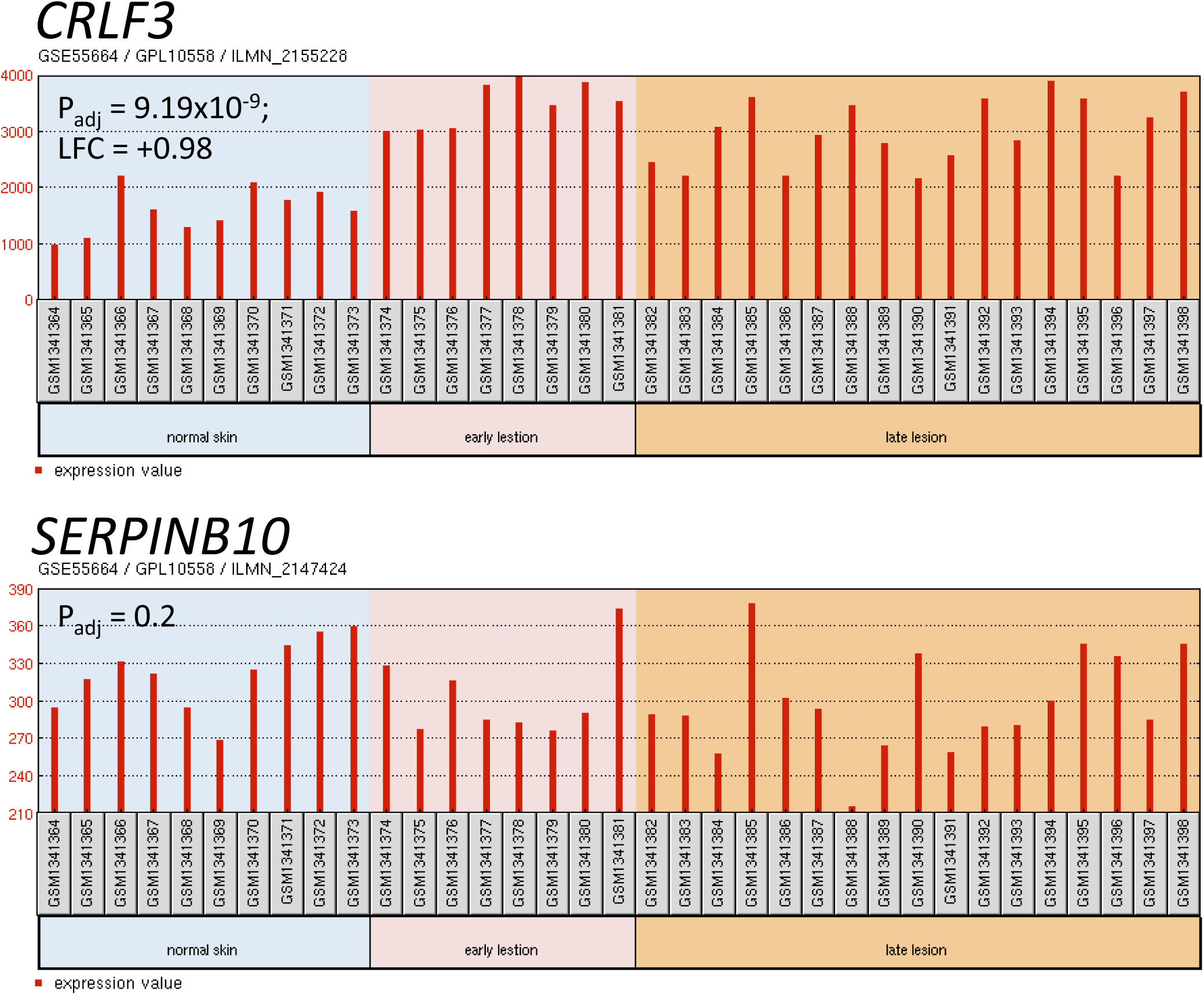
Expression of novel candidate genes derived from the GWAS data in RNA from biopsies of normal skin from healthy donors compared to biopsies from lesions of CL patients. Data were accessed from the GEO database: GSE55664.^1^ Between group comparisons (normal versus lesions) were made on log transformed data using the GEO2R tool within the GEO database, with P-values adjusted using the Benjamini and Hochberg false discovery rate.

**Table S1.** Characteristics of the Post-QC Phase 1 and Phase 2 Samples.

|  | Phase 1 |  |  | Phase 2 |  |  |
| --- | --- | --- | --- | --- | --- | --- |
|  | CL cases | Endemic Controls | Blood Bank Controls | CL cases | Endemic Controls | Blood Bank Controls |
| N° participants | 956 | 237 | 631 | 1110 | 604 | 574 |
| Males | 560 | 89 | 420 | 659 | 273 | 346 |
| Females | 396 | 148 | 211 | 451 | 331 | 228 |
| Age at collection |  |  |  |  |  |  |
| Male Mean±SD | 29±16 | 24±19 | 35±11 | 31±16 | 18±13 | 37±10 |
| Male Range | 1-78 | 2-81 | 17-65 | 5-87 | 2-75 | 16-65 |
| Female Mean±SD | 28±15 | 30±19 | 32±10 | 32±16 | 22±15 | 35±10 |
| Female Range | 4-75 | 1-88 | 16-68 | 3-85 | 2-72 | 17-65 |

**Table S2.** Signals of Association at Candidate and Related Loci Previously Reported as Genetic Risk Factors for CL Caused by *L. braziliensis*.

| Chr | Position (bp) | rsID | P-value | Odds Ratio (95% CI) | Beta (SE) | Allele<sup>1</sup> | Variant Origin | Location | Gene Symbol<sup>2</sup> | Function (Origin of Variant) |
| --- | --- | --- | --- | --- | --- | --- | --- | --- | --- | --- |
| 1 | 86264025 | rs380654 | 2.06E-04 | 0.95 (0.93-0.98) | -0.29 (0.008) | A (A/C) | Global | Intron | COL24A1 | Collagen family member 24A1 |
| 1 | 92303128 | rs141665520 | 3.98E-04 | 1.15 (1.07-1.25) | 0.026 (0.007) | C (G/C) | African | intron | TGFBR3 | TGF-β receptor 3 |
| 1 | 92306871 | rs77138249 | 3.98E-04 | 1.15 (1.07-1.25) | 0.026 (0.007) | C (T/C) | African | intron | TGFBR3 | TGF-β receptor 3 |
| 1 | 103417667 | rs114806195 | 6.22E-04 | 0.90 (0.85-0.96) | -0.025 (0.007) | T (T/A) | Global | intron | COL11A1 | Collagen family member 11A1 |
| 1 | 103418632 | rs111420962 | 6.22E-04 | 0.90 (0.85-0.96) | -0.025 (0.007) | T (A/T) | Global | intron | COL11A1 | Collagen family member 11A1 |
| 1 | 154428283 | rs12133641 | 7.14E-04 | 0.96 (0.94-0.98) | -0.027 (0.008) | T (A/T) | Global | intron | IL6R | Interleukin 6 receptor |
| 4 | 146477459 | rs76796874 | 7.49E-04 | 1.15 (1.06-1.24) | 0.026 (0.008) | A (G/A) | African | intron | SMAD1 | R-SMAD<sup>3</sup> |
| 13 | 37445256 | rs8001427 | 0.004 | 1.13 (1.04-1.23) | 0.022 (0.008) | T (A/T) | African | Intron | SMAD9 | R-SMAD<sup>3</sup> |
| 15 | 67079073 | rs59361088 | 0.001 | 0.86 (0.79-0.94) | -0.024 (0.007) | A (C/A) | African | downstream | SMAD6 | I-SMAD<sup>3</sup> |
| 15 | 67364991 | rs4776880 | 0.009 | 0.96 (0.93-0.99) | -0.021 (0.008) | T (G/T) | Global | intron | SMAD3 | R-SMAD<sup>3</sup> |
| 18 | 45390603 | rs115582038 | 1.47E-04 | 1.19 (1.09-1.30) | 0.028 (0.007) | C (G/C) | African | intron | SMAD2 | R-SMAD<sup>3</sup> |
| 18 | 45393852 | rs75753347 | 1.47E-04 | 1.19 (1.09-1.30) | 0.028 (0.007) | T (A/T) | African | intron | SMAD2 | R-SMAD<sup>3</sup> |
| 18 | 46454258 | rs9956511 | 0.005 | 0.86 (0.77-0.95) | -0.021 (0.008) | C (A/C) | African | intron | SMAD7 | I-SMAD<sup>3</sup> |
| 18 | 48562852 | rs78801230 | 6.02E-04 | 1.15 (1.06-1.25) | 0.026 (0.008) | T (A/T) | European | intron | SMAD4 | Co-SMAD<sup>3</sup> |
| 18 | 48604010 | rs1241993461 | 1.90E-04 | 1.17 (1.08-1.27) | 0.028 (0.008) | G (A/G) | Rare | intron | SMAD4 | Co-SMAD<sup>3</sup> |
| 21 | 34654499 | rs2247177 | 0.003 | 0.97 (0.95-0.99) | -0.023 (0.008) | A (G/A) | Global | intron | IL10RB | Interleukin 10 receptor B |

**NOTE** <sup>1</sup>Associated allele (ancestral/minor) for risk or protection as indicated by the odds ratio. <sup>2</sup>Data for two variants provided when the top two hits were in linkage disequilibrium. <sup>3</sup>SMADs are structurally similar proteins that are the main signal transducers for receptors of the transforming growth factor beta (TGF- $\beta$ ) superfamily. The abbreviation derives from homologies to the *Caenorhabditis elegans* SMA ("small" worm phenotype) and *Drosophila* MAD ("Mothers Against Decapentaplegic") family of genes. R-SMAD = receptor-regulated SMAD; Co-SMAD = common partner SMAD; I-SMAD = inhibitory SMAD.

## Footnotes

1 Pruim RJ, Welch RP, Sanna S, et al. LocusZoom: regional visualization of genome-wide association scan results. Bioinformatics **2010**; 26:2336-7.

1 Novais FO, Carvalho LP, Passos S, et al. Genomic profiling of human Leishmania braziliensis lesions identifies transcriptional modules associated with cutaneous immunopathology. J Invest Dermatol **2015**; 135:94-101.

